# DNA Self-Assembled Plasmonic Nanodiamonds for Biological Sensing

**DOI:** 10.1101/2021.11.09.467982

**Authors:** Le Liang, Peng Zheng, Sisi Jia, Krishanu Ray, Yun Chen, Ishan Barman

## Abstract

Nitrogen-vacancy (NV) centers in diamonds are promising solid-state quantum emitters for developing superior biological imaging modalities. They possess desired bio-compatibility, photostability and electronic spin-related photophysical properties that are optically accessible at room temperature. Yet, bare nanodiamond-based imaging modalities are limited by the brightness and temporal resolution due to the intrinsically long lifetime of NV centers. Moreover, it remains a technological challenge using top-down fabrication to create freestanding hybrid nanodiamond imaging probes with enhanced performance. In this study, we leverage the bottom-up DNA self-assembly to develop a hybrid plasmonic nanodiamond construct, which we coin as the **p**lasmon-**e**nhanced **n**anodiamond (PEN), for biological imaging. The PEN nano-assembly features a closed plasmonic nanocavity that completely encapsulates a single nanodiamond, thus enabling the largest possible plasmonic enhancement to accelerate the emission dynamics of NV centers. Creation of the PEN nano-assembly is size-independent, so is its broadband scattering spectrum that is optimally overlapped with the emission spectrum of NV centers. Study of the structure-property correlation reveals that the optimal condition for emission dynamics modification is causally linked to that for a plasmonic nanocavity. The cellular internalization and cytotoxicity studies further confirm the delivery efficiency and biological safety of PEN nano-assemblies. Collectively, the PEN nano-assembly provides a promising approach for manipulating photophysical properties of solid-state quantum emitters and could serve as a versatile platform to uncover non-trivial quantum effects in biological systems.

## 1. Introduction

Nitrogen-vacancy (NV) centers in diamonds have recently emerged as a unique type of solid-state quantum emitters. They have found an array of exciting applications in biological imaging, metrology, electric and magnetic field sensing, and quantum information technology, owing to their bio-compatibility, photostability and electronic spin-related photophysical properties that are optically accessible at room temperature.^1-5^ The prospect of unlocking the potential of NV centers for developing superior solid-state optical devices hinges on our ability to control and harness their distinct photophysical properties at the single quantum emitter level. Localized surface plasmon resonance (LSPR) possesses a highly confined local electromagnetic (EM) field at the subwavelength scale, broadband plasmonic resonance, ultrafast dynamics, and extraordinary spatial coherence,^6-7^ and therefore they provide a promising route to manipulate the emission dynamics of a single NV center. However, the EM field of LSPR decays rapidly over a short distance (∼10 nm).^8-9^ Further exacerbating the issue is the finite size of a nanodiamond varying from a few to hundreds of nanometers that makes plasmon-NV centers inefficiently coupled.^10-11^ Such a length-scale incompatibility remains a significant unmet technological challenge and handicaps current efforts towards developing a versatile nanophotonic platform to maximize interactions between LSPR and NV centers.

To effectively augment the plasmonic coupling and mitigate the size effect, the concept of plasmonic nanocavities has been explored.^12^ Underpinning a plasmonic nanocavity is the amplified local density of optical states (LDOS) enclosed,^13-15^ which is crucial to accelerate the photon emission rate and increase the quantum emitter brightness for creating superior imaging modalities with enhanced temporal resolution and sensitivity.^16-18^ While open nanocavities are readily achievable using top-down lithography methods,^19-20^ their performance is compromised by the modest EM field confinement. Instead, closed nanocavities are desired as they can deliver an intensely concentrated EM field for efficient near-field plasmonic interaction with the enclosed quantum emitter.^17^ Recent progresses on creating semi-closed nanocavities have demonstrated impressive lifetime shortening and brightness enhancement for NV centers positioned within, including nanoparticle dimers^21-22^ and nanopatch antennas^23-25^. Nevertheless, these semi-closed nanocavities prove inept to deliver a spatially uniform LDOS and are limited to efficiently interact with only small nanodiamonds (typically less than 35 nm), owing to the rapid spatial decay of the plasmonic field.^26-27^ Moreover, the dimer and nanopatch antenna constructs are difficult to be translated into free-standing imaging modalities that also demand size tunability for optimal cellular internalization in biological applications. The promise and challenges associated with these semi-closed nanocavities highlight the need to innovate synthetic strategies for creating closed nanocavities, which could hold the key to deliver superior plasmonic nanodiamonds-based imaging modalities.

Recently, DNA self-assembly has proved a powerful bottom-up approach. As compared to the top-down lithography methods that require a mask to pre-define the resulting structure, DNA self-assembly is a highly versatile and programmable synthetic approach and enables the formation of a wide variety of pre-defined constructs through spontaneous self-ordering and organization of DNA structural constituents that are driven by DNA hybridization. ^28-31^ Success of DNA self-assembly has been manifested as a wide array of preciously constructed DNA metamolecules,^6, 32-36^ which possess exciting new photophysical properties rarely seen on an individual constituent. Further underscoring the promise of DNA self-assembly is our own recent breakthrough on developing the DNA-STROBE method that has been exploited for single-molecule plasmonic sensing.^37^ Given that colloidal nanodiamonds and gold nanoparticles can be readily modified with single-stranded DNA (ssDNA) sequences, they thus can serve as ideal DNA structural constituents and spontaneously re-organize into the pre-defined nanoscale constructs, including the highly desired closed plasmonic nanocavity with a nanodiamond encapsulated within, through DNA hybridizations.

In this study, by leveraging the versatility and programmability of DNA self-assembly, we report a hybrid plasmonic nanodiamond construct, which we coin as **p**lasmon-**e**nhanced **n**anodiamond (PEN). The PEN nano-assembly features a closed plasmonic nanocavity formed by a cluster of gold nanoparticles that completely encapsulates a single nanodiamond in a core-satellite manner. Creation of the PEN nano-assembly is size-independent, so is its broadband scattering spectrum that is optimally overlapped with the emission spectrum of NV centers. Combined with the spatial proximity of the core-satellite construct, the PEN nano-assembly facilitates the largest possible interaction between the cavity mode and the enclosed NV centers. Correlative measurements using scanning electron microscopy (SEM), dark-field microscopy, and fluorescence lifetime imaging microscopy (FLIM), not only unequivocally demonstrate the emission rate enhancement for the enclosed NV centers, but also uncover a causal relationship between the optimal condition for emission dynamics modification and that for a plasmonic nanocavity, which further underscores the role of a plasmonic nanocavity in modifying the photophysical properties of the enclosed quantum emitters. Further studies on the interplay of PEN nano-assemblies and HeLa cells confirm the internalization efficiency and biological safety of the constructs. In addition to enhancing the emission dynamics of NV centers, the PEN nano-assembly can be readily extendable for monitoring dynamic intracellular pH^38^ or temperature^39-40^ and serve as solid-state single-photon emitters.

## 2. Results and Discussion

### 2.1. Photophysical properties of the PEN nano-assembly as compared to the dimer construct

The PEN nano-assembly supports a plasmonic nanocavity that completely encloses a nanodiamond, which is in sharp contrast to the semi-open nanocavity of the gold-nanodiamond-gold dimer construct. To gain a mechanistic understanding of the two types of nanocavities with different sophistications, we leveraged finite-difference time-domain (FDTD) simulations to study their distinct photophysical properties. In FDTD simulations, the PEN nano-assembly was modelled to consist of a spherical nanodiamond as the core surrounded by a cluster of gold nanoparticles, as schematically shown and compared to the dimer in Fig. 1a. All the nanoparticles were modeled to have a diameter of 40 nm. A single NV center was assumed in the nanodiamond and modelled as an electric dipole oscillating horizontally with an intrinsic quantum efficiency of 0.7.

**Figure 1.**
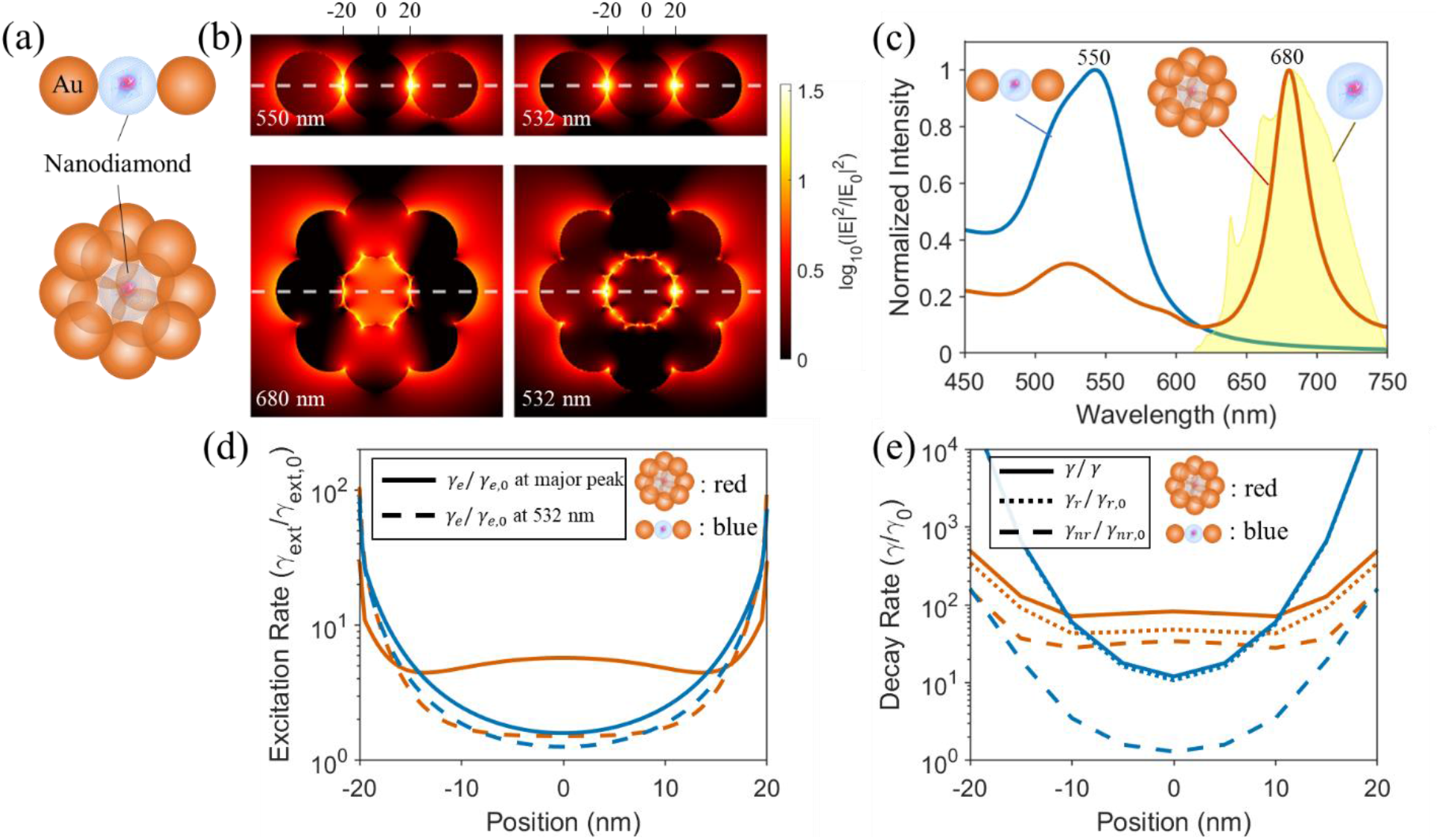
Comparison of photophysical properties of a gold-nanodiamond-gold dimer and the PEN nano-assembly. (a) Schematics for the dimer and PEN constructs. All the modelled nanoparticles have the same diameter of 40 nm. (b) Calculated electric field enhancement for both constructs at 532 nm and their major resonance peak wavelengths as specified in (c). A horizontal incident polarization was used. (c) Calculated extinction spectra for both constructs as compared to the experimentally measured fluorescence emission spectrum for nanodiamonds (yellow shaded) with a 532 nm excitation. (d) The line prolife as indicated by the white dashed line in (b) of the calculated excitation rate enhancement for both constructs at 532 nm and their major resonance peak wavelengths. (e) The line prolife as indicated by the white dashed line in (b) of the calculated enhancement for total decay rate (solid curve), radiative decay rate (short-dashed curve), and nonradiative decay rate (dashed curve) for both constructs at their major resonance peak. All the calculation were done using FDTD simulations, assuming an intrinsic quantum efficiency of 0.7 for the used electric dipole oscillating horizontally.

From FDTD simulations, the most intense hotspots were formed at the gold-nanodiamond interface for both constructs (Fig. 1b). Because of the large gap defined by the nanodiamond, the dimer displayed a weak plasmonic coupling, which was also reflected on the barely deconvoluted extinction peaks in Fig. 1c. In comparison, as the surrounding gold nanoparticles of the PEN nano-assembly formed a closed nanocavity, an intense electric field enhancement was produced and uniformly distributed over the entire nanocavity. Importantly, the plasmonic nanocavity mode of the PEN nano-assembly has a strong spectral overlap with the emission spectrum of NV centers (yellow shaded in Fig. 1c), which is a prerequisite for intense plasmon-NV center interactions. In contrast, such a spectral overlap is difficult to achieve for the dimer, as a smaller gap (less than 10 nm) is required for its plasmonic resonance to be significantly redshifted.

The excitation and decay rate enhancements have been calculated as a function of the NV center position within the nanocavity. From Fig. 1d-e, the PEN nano-assembly displayed a much larger and almost position-independent excitation and decay rate enhancements (including radiative, nonradiative, and total decay) inside the nanocavity. This is in sharp contrast to the dimer, in which both excitation and decay rate enhancements dropped significantly when the NV center moved from the nanodiamond edge to the center. Although the dimer showed a considerable total decay rate enhancement, it was dominated by the nonradiative component (shorted dashed curve in Fig. 1e), thus contributing little to photon yield. The above observations underscore advantages of using a closed plasmonic nanocavity for emission dynamics modification as well as the unexplored promise of the PEN nano-assembly for developing desired imaging modalities with enhanced photon emission rate.

### 2.2. Fabrication of PEN nano-assemblies based on DNA self-assembly

To fabricate the PEN nano-assembly, we leveraged DNA self-assembly, Previously, we have successfully employed DNA self-assembly to develop a hierarchical core-satellite nano-assembly for single-molecule plasmonic sensing.^37^ Following the same method, we started to prototype a PEN nano-assembly by modifying biotin-capped nanodiamonds (diameter ∼100 nm) and bare gold nanoparticles (diameter ∼50 nm) using two complementary ssDNA sequences. By mixing ssDNA-modified nanodiamonds and gold nanoparticles with the latter in excess, PEN particles of the desired configuration were achieved. The fabrication protocol is schematically shown in Fig. 2a. Successful fabrication of the PEN nano-assemblies was validated by SEM imaging of nanodiamonds before and after DNA self-assembly, as shown in Fig. 2b. Both biotin-capped nanodiamonds and the fabricated PEN nano-assemblies displayed excellent dispersibility without noticeable aggregation, highlighting the robustness and reproducibility of this synthetic approach. Expanding on this demonstration, we have further fabricated PEN nano-assemblies with a variety of size combinations for nanodiamonds and gold nanoparticles, as shown in Fig. 3, demonstrating the versatility and size independence of this method.

**Figure 2.**
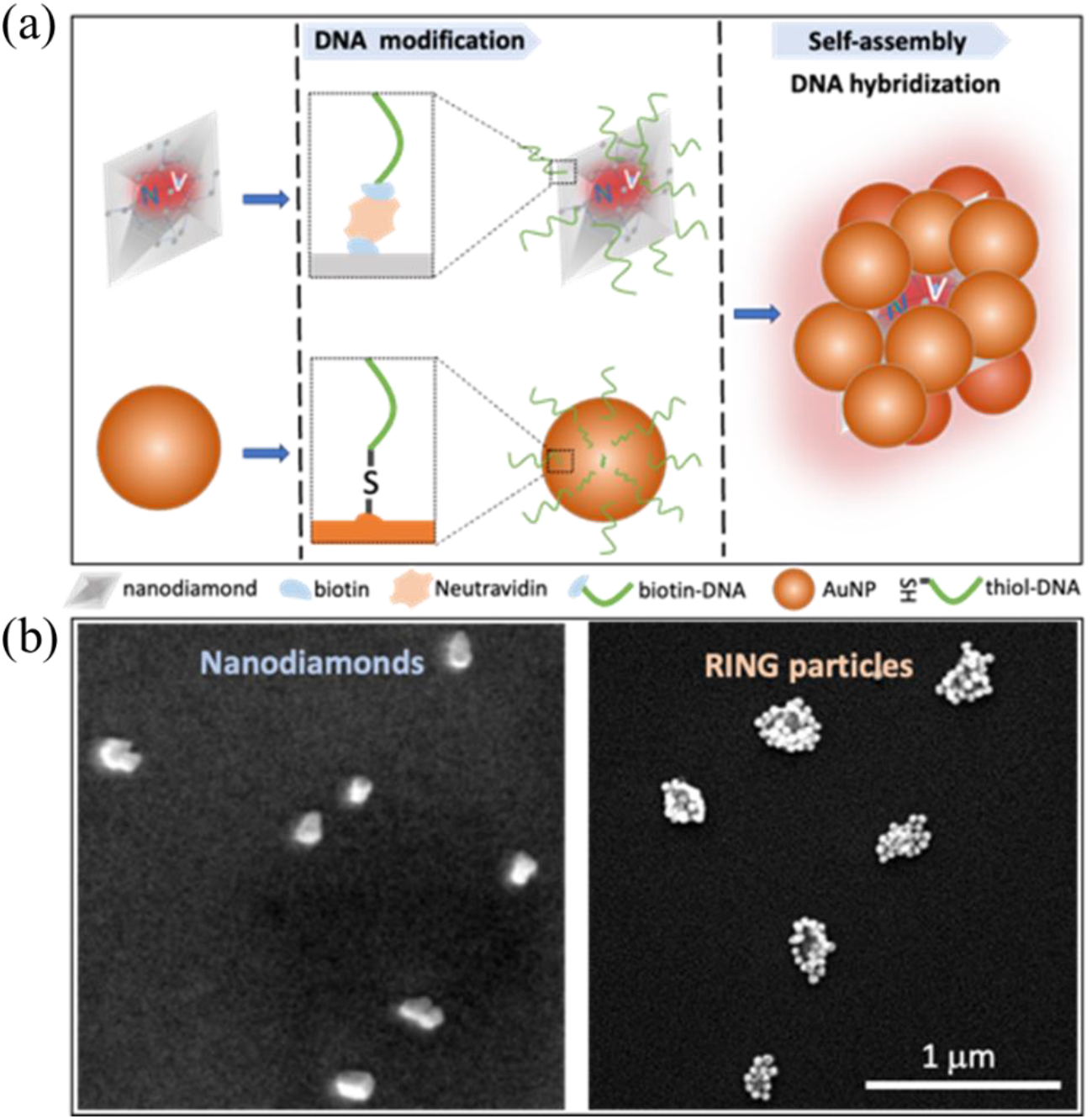
An overview of the synthetic route for PEN nano-assemblies by DNA self-assembly. (a) Biotin-capped nanodiamonds (diameter: 100 nm) and gold nanoparticles (diameter 50 nm) are modified by complementary single-stranded DNA sequences, and upon DNA hybridization, PEN nano-assembly is formed. (b) SEM images of representative biotin-capped nanodiamonds (left) and the synthesized PEN nano-assemblies (right), featuring excellent dispersion without noticeable aggregation.

**Figure 3.**
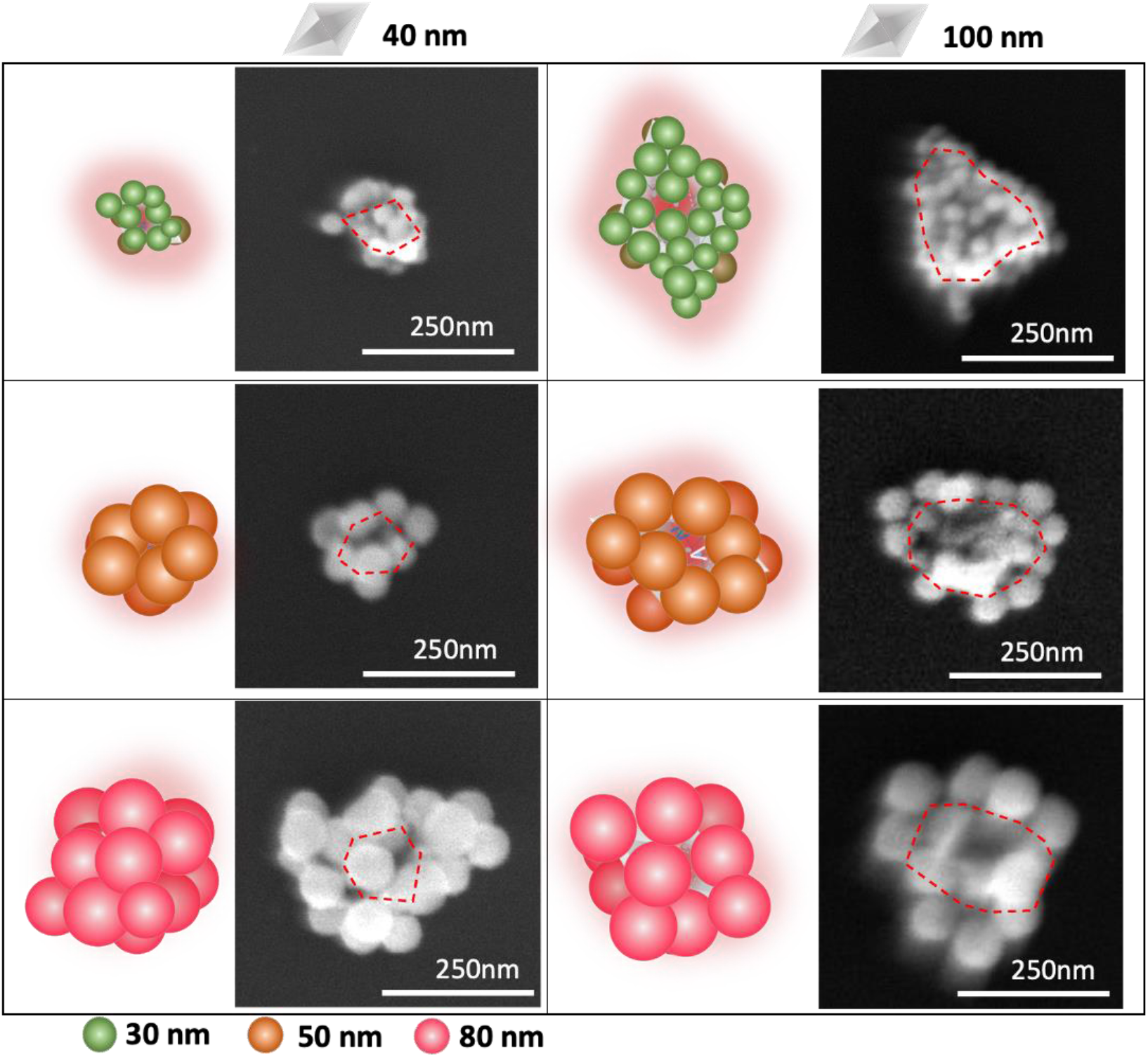
PEN nano-assemblies with different size combinations of nanodiamonds (diameter: 40 nm, 100 nm) and gold nanoparticles (diameter: 30 nm, 50 nm, 80 nm) constructed by DNA self-assembly. Successful fabrication of different variants of PEN nano-assemblies underscores the versatility of the synthetic method. First and third column: schematics; second and fourth column: SEM images.

### 2.3. Photophysical properties of the PEN nano-assembly

To understand how the emission dynamics of NV centers in a nanodiamond is modified by plasmonic effects, we performed correlative photophysical studies for a representative PEN nano-assembly made of a nanodiamond of 100 nm and gold nanoparticles of 80 nm in diameter, as shown in Fig. 4. We designed special fiducial markers on indium tin oxide (ITO)-coated glasses for locating the PEN nano-assembly across different modalities (Fig. 4a). The studied PEN nano-assembly, as indicated by the white arrow, was clearly seen under SEM (Fig. 4b). The same PEN nano-assembly was successively characterized by the dark-field microscopy and FLIM. The dark-field scattering spectrum featured a broadband plasmonic resonance between 550 nm to 750 nm (Fig. 4c), which is typical of a cluster of gold nanoparticles owing to the overall large dimension and interparticle coupling.^41^ The corresponding fluorescence spectrum displayed a broad emission band between 630 nm to 775 nm, which is characteristic of the spontaneous emission from bare nanodiamonds (Fig. 4d). The zero-phonon line at about 638 nm, which is attributed to the negatively charged (NV^-^) state of the nitrogen vacancy center,^21^ was also well discernible. The FLIM image showed a mean lifetime of 2∼3 ns for the studied PEN particle, which is a considerable reduction from a typical lifetime of 13 ns at the NV center in bare nanodiamonds (intrinsic lifetime information provided by the vendor). The lifetime shortening is a clear manifestation of the accelerated emission dynamics of NV centers because of the newly created LDOS within the enclosed plasmonic nanocavity. Indeed, the strong scattering and emission spectral overlap of the PEN nano-assembly, combined with the spatial proximity of the core-satellite construct, significantly facilitates intense coupling of the plasmon-NV centers. Such experimental observations also corroborate our numerical analysis on the advantages of using a closed plasmonic nanocavity for modifying the emission dynamics of a quantum emitter in Fig. 1.

**Figure 4.**
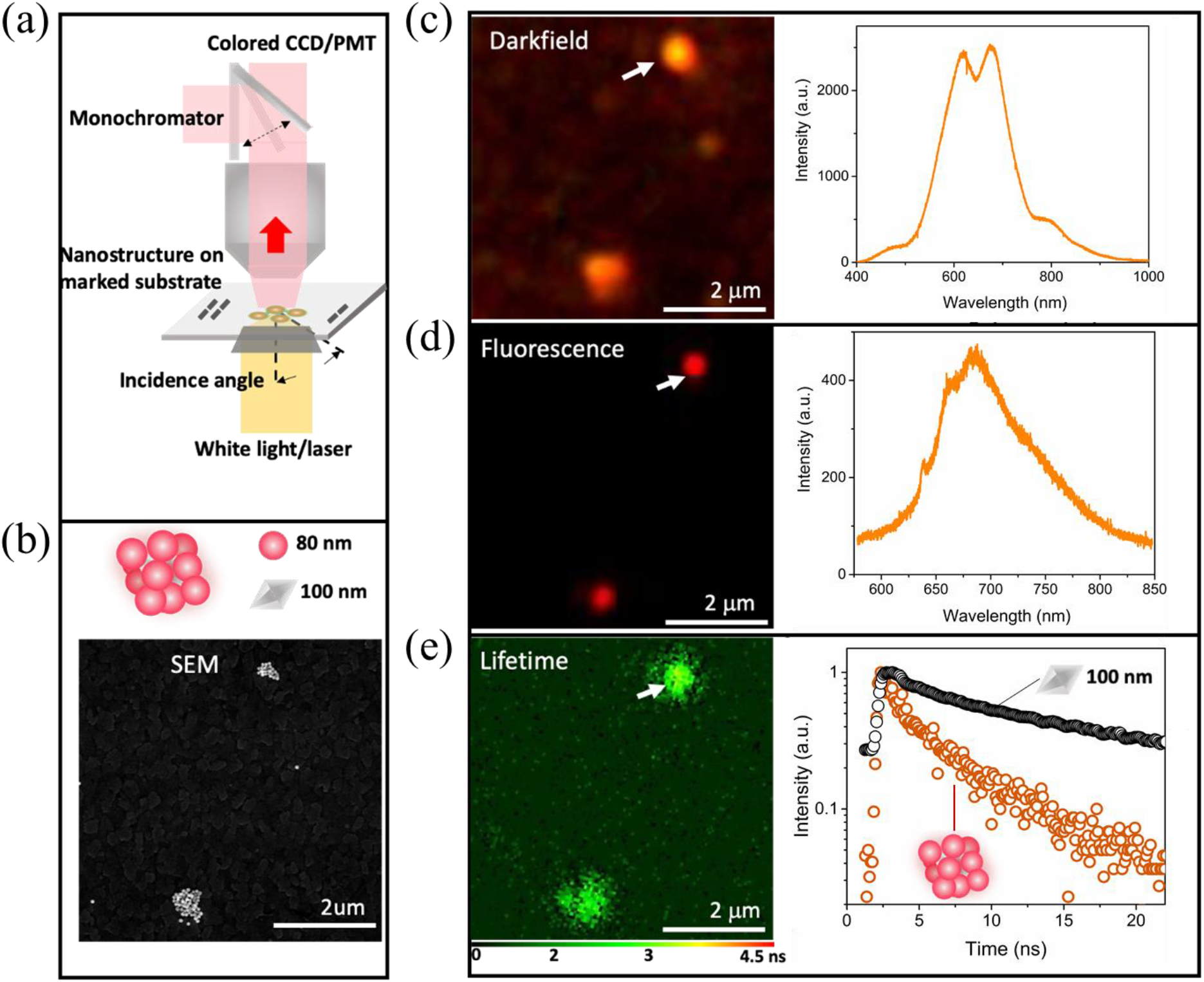
Correlative characterizations for PEN nano-assemblies using SEM, dark-field microscopy and FLIM. (a) Schematic of the imaging system setup. (b) SEM image, (c) dark-field image and scattering spectrum, (d) fluorescence image and emission spectrum, and (e) FLIM image and lifetime decays for the studied PEN nano-assembly as indicated by the white arrow.

### 2.4. Structure-property relationship of PEN nano-assemblies

Given the wide parameter tunability of the PEN nano-assembly, we strive to uncover its structure-property relationship by a systematic investigation of different variants of the construct. To this end, we expanded correlative photophysical characterizations to all the fabricated PEN nano-assemblies with a variety of size combinations (Fig. 3). The obtained results using SEM, dark-field, and FLIM images for each parameter combination were displayed in Fig. 5. For all the studied PEN nano-assemblies, despite dramatically different size combinations, they ubiquitously displayed a broadband scattering peak between 600 nm and 800 nm, resulting in an optimal spectral overlap with the emission spectrum of NV centers. The emergence of subpeaks for large PEN nano-assemblies seen in Fig. 5e was likely contributed by higher-order plasmonic modes which are typical for plasmonic nanoparticles with a larger dimension.^42-43^

**Figure 5.**
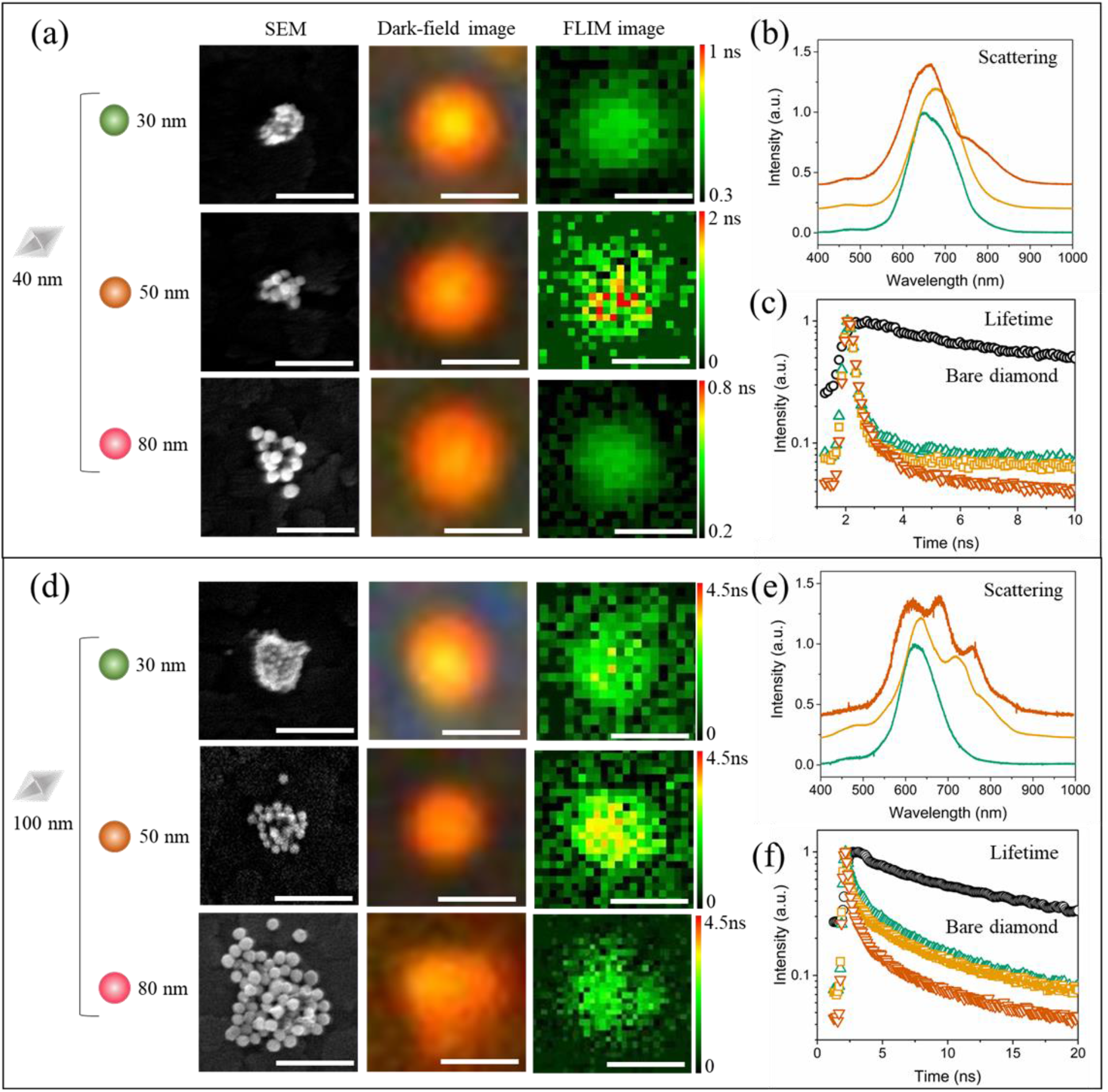
Correlative studies for PEN nano-assemblies with different size combinations. (a, d) SEM (first column), dark-field (second column), and FLIM (third column) for PEN nano-assemblies with the parameters as specified on the left. (b, e) Scattering spectra extracted from dark-field images. (c, f) Average lifetime decays from the FLIM images. For (b, c, e, f), the blue, orange and red curves/symbols represent PEN nano-assemblies made of gold nanoparticles with a size of 30 nm, 50 nm, and 80 nm, in consistent with the symbols used on the left of (a, d). The scale bars for SEM, dark-field image, and FLIM are 500 nm, 1 µm, and 1 µm.

Different from the largely size-independent scattering spectra, the emission dynamics of NV centers confined in the PEN nano-assembly is size-dependent. Globally, the PEN nano-assemblies enclosing a smaller nanodiamond exhibit a faster emission dynamics (Fig. 5c, f). A close examination revealed that, for the same nanodiamond enclosed by larger gold nanoparticles, or for a smaller nanodiamond enclosed by the same gold nanoparticles, NV centers experience a faster decay dynamics. These observations suggest that the PEN nano-assembly enables an enhanced emission rate for NV centers, and more importantly, the emission rate can be further augmented if a smaller nanodiamond and/or larger gold nanoparticles are used. Such an optimal condition for the emission dynamics modification is not merely a coincidence with the requirement for an optimized plasmonic nanocavity, it further reinforces the role the plasmonic nanocavity plays in modulating the emission dynamics of the enclosed NV centers. Our finding underpins the distinct plasmonic attributes of the PEN nano-assembly for enhancing photophysical properties of the enclosed quantum emitters.

Additionally, revelation of the structure-property relationship has profound implications in applying PEN nano-assemblies to image intracellular processes. As cells internalize nano- or micro-sized objects via various size-dependent mechanisms, including phagocytosis, clathrin-mediated endocytosis, and caveolin-mediated endocytosis, the size of PEN nano-assemblies can be customized to target specific pathway. Because neither the synthetic approach for creating, or the scattering spectrum of the PEN nano-assembly, is size-dependent, the size customization can be performed without significant impact on photon yields.

### 2.5. Quantification of the optimal emission dynamics modification

The uncovered structure-property relationship suggested that the largest possible modification to the emission dynamics could be achieved on a PEN nano-assembly made of the smallest nanodiamond and the largest gold nanoparticles. Therefore, we fabricated PEN nano-assembly using nanodiamonds of 40 nm and gold nanoparticles of 80 nm to optimize the emission rate and brightness. To gain a quantitative picture of the extent to which the emission dynamics of NV centers was modified, a statistical analysis was performed on the well-dispersed biotin-capped nanodiamonds of 40 nm and the corresponding PEN nano-assemblies with a cluster of gold nanoparticles of 80 nm. It was observed that the mean lifetime of NV centers decreased from 7.94 ns in bare nanodiamonds on ITO-coated glass to 0.45 ns after the nanodiamonds were enclosed into the PEN nano-assembly on the same substrate (Fig. 6a). In other words, the increased plasmonic LDOS inside the nanocavity resulted in 18-fold emission rate enhancement. Moreover, the same NV centers were found to exhibit 50-fold fluorescence intensity enhancement (Fig. 6b), which is a combined effect of excitation rate enhancement and quantum efficiency enhancement. The amplified emission rate and brightness of NV centers underscore the exciting potential of exploiting PEN nano-assemblies to develop superior sensors for cellular imaging.

**Figure 6.**
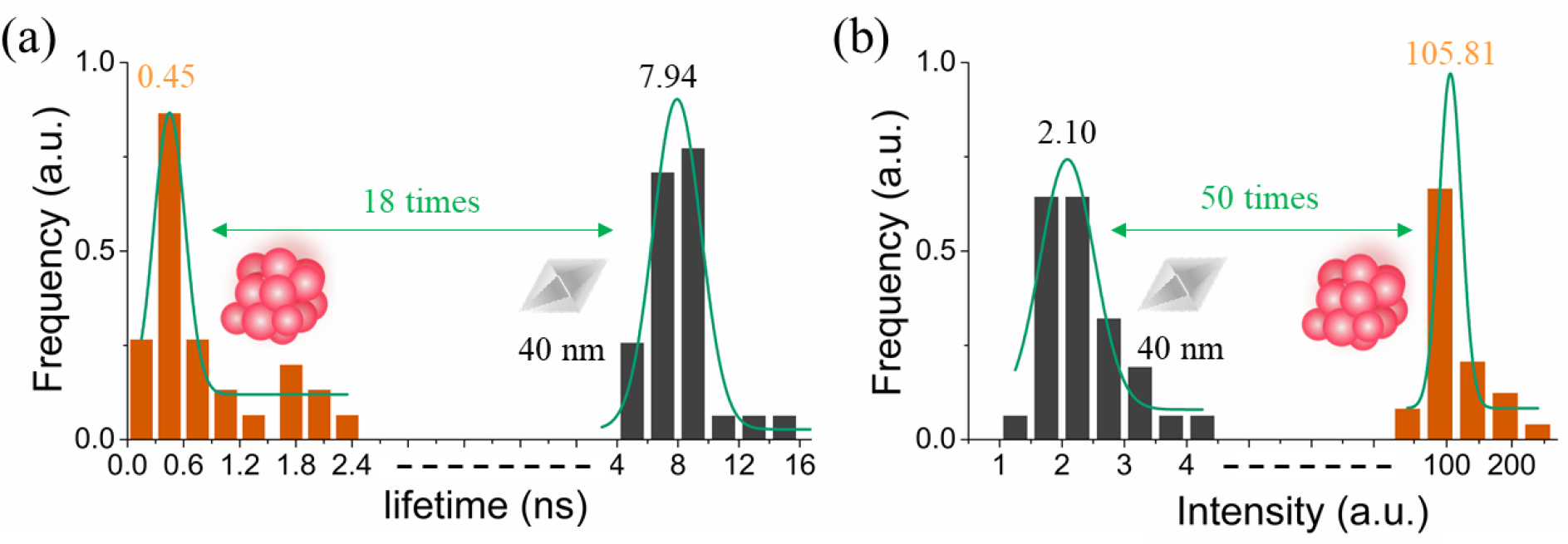
Statistical analysis of (a) the lifetime and (b) the emission intensity for the PEN nano-assembly made of 40 nm nanodiamond and 80 nm gold nanoparticles as compared to these of bare nanodiamonds with the same size.

### 2.6. Cellular internalization and cytotoxicity of PEN nano-assemblies

To evaluate the feasibility of applying the PEN nano-assembly for cellular imaging, we studied its internalization efficiency and cytotoxicity. The PEN nano-assemblies were incubated with GFP-labelled HeLa cells at 37 °C while the uptake of PEN was monitored by confocal fluorescence microscopy for 12 hours. The 2D and 3D images confirmed that PEN nano-assemblies were internalized by cells. After the internalization at 3 hours post-incubation, we observed that PEN nano-assemblies were further transported within the cell. A concentrated distribution was detected near the nuclei after 12 hours of incubation. To study the cytotoxicity, the PEN nano-assemblies were incubated with Hela cells for 3 days. The ethidium homodimer was subsequently used to stain dead cells. We found that cell viability of 98.9% ± 0.75% of the cells incubated with PEN nano-assemblies was statistically indistinguishable from the control group (Fig. 7b). We concluded that PEN nano-assemblies are suitable for cellular imaging given their high cell-permeability and low cytotoxicity.

**Figure 7.**
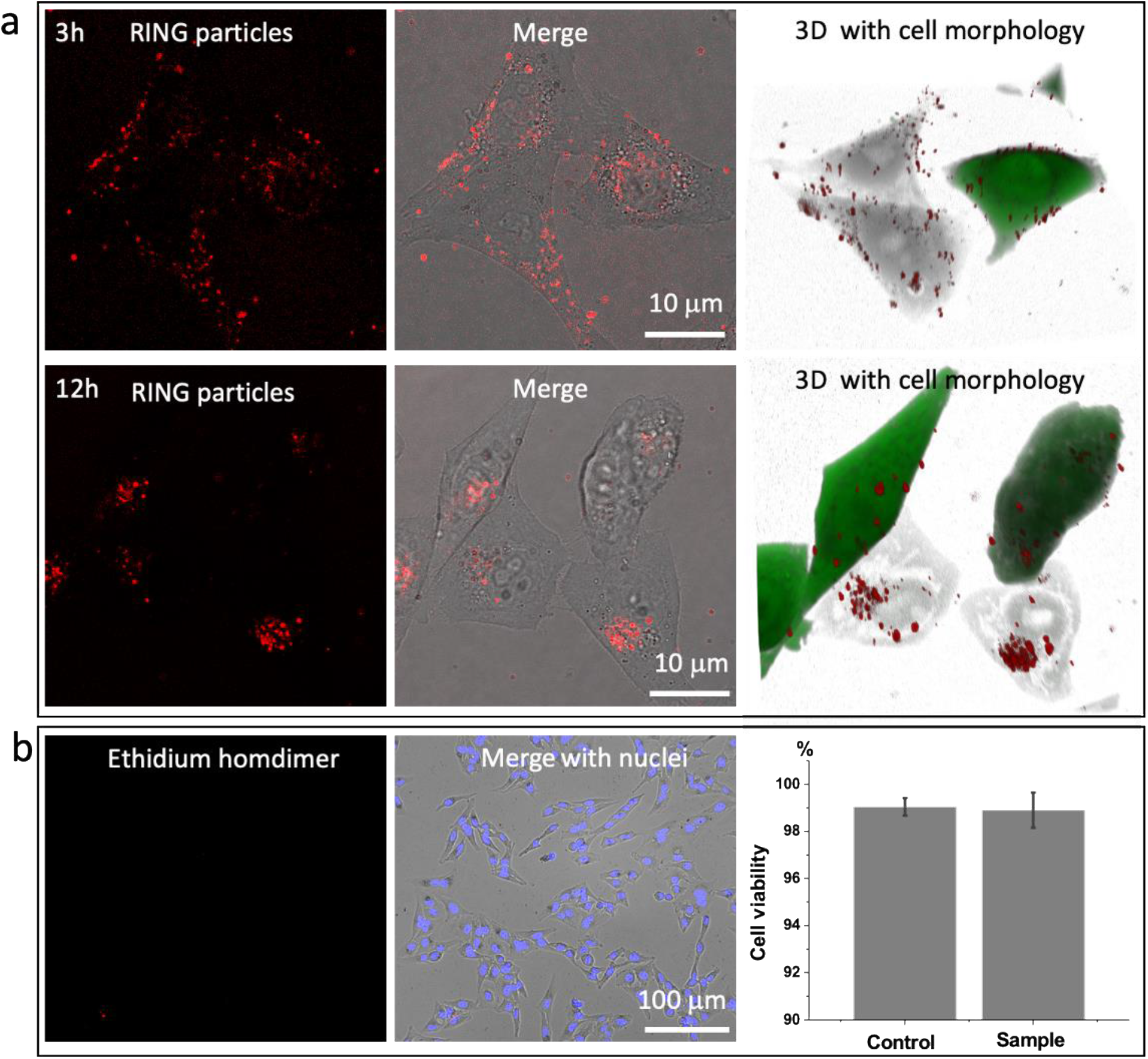
Evaluation of PEN nano-assemblies in cells. (a) Confocal fluorescence images of PEN nano-assemblies (red) inside cells with 3D reconstruction after 3 hours’ and 12 hours’ incubation. (b) Cytotoxicity assessment of PEN nano-assemblies by confocal fluorescence images. Dead cells were stained by ethidium homodimer (red) after 3 days incubation. The cell viability was found to be 98.9% ± 0.75%.

## 3. Conclusion

In this study, we fabricated a PEN nano-assembly to modulate the emission dynamics of NV centers in nanodiamonds. The PEN nano-assembly can strongly accelerate the emission rate of NV centers owing to the closed plasmonic nanocavity that supports a large LDOS. The versatility of DNA self-assembly also allows creation of numerous variants of PEN nano-assemblies with different size combinations of nanodiamonds and gold nanoparticles. A systematic investigation of the structure-property relationship uncovers that the optimal condition for emission dynamics modification is causally linked to that for a plasmonic nanocavity, validating the superiority of a closed plasmonic nanocavity over an open or semi-open nanocavity. Furthermore, the PEN nano-assemblies display excellent cell permeability and negligible cytotoxicity. In summary, PEN nano-assemblies provide a robust, versatile, and bio-compatible approach to realize enhanced emission rate and brightness in nanodiamonds with NV centers, which could pave the way for studying a wide variety of dynamic biological processes.

### Materials and methods

#### Preparation of gold nanoparticles and oligo DNA conjugation

Prior to use, the colloidal solutions of 30 nm, 50 nm and 80 nm AuNPs (BBI TM Solutions) were subjected to centrifugation (30 nm AuNPs: 10000 rpm for 10 min; 50 nm AuNPs: 6000 rpm for 10 min; 80 nm AuNPs: 4000 rpm for 10 min). The disulfide bond in the thiol-modified oligonucleotides was reduced to monothiol using TCEP (20 mM, 1h). The oligonucleotides were purified using size exclusion columns (G-25, GE Healthcare) to get rid of the small molecules. 800 μL of the concentrated colloidal solution of 80 nm AuNPs was mixed with freshly dissolved thiol-modified DNA (100 μM) in a 1:50,000 ratio (1:10,000 for 30 nm AuNPs and 1:20,000 for 50 nm AuNPs) in water, and the mixture was incubated for 2 hours at room temperature (300 rpm). 100 μL PB buffer (100 mM, pH=7.4) was then added to the mixture. After 30 min, we added 10 μL of NaCl solution (2 M) every 20 min for 4 times, and then 20 μL NaCl solution (2 M) every 30 min for 3 times. The NaCl concentration was gradually increased to ensure the full coverage of AuNPs with thiolated DNA. The final concentration of NaCl was 200 mM, and the mixture was incubated at room temperature (300 rpm) overnight. The AuNP-DNA conjugates were purified by centrifugation. Freshly prepared, fully covered 30 nm, 50 nm and 80 nm AuNPs did not precipitate in the PBS buffer (100 mM, pH 8.0). This high salt resistance property of fully covered AuNPs makes it possible to assemble the core-satellite structures through DNA hybridization.

#### Preparation of diamond nanoparticles and oligo DNA conjugation

Prior to use, the colloidal solutions of 40 nm and 100 nm biotin-modified nanodiamond (1m/ml Adámas Nanotechnologies) were mixed with freshly dissolved neutravidin (10µg/ml) in a 1:100 ratio v/v (1:10 for 100 nm nanodiamond) in water, and the mixture was incubated for 2 hours at room temperature (300 rpm). The nanodiamond-avidin conjugates were purified by centrifugation (40 nm nanodiamond: 5000 rpm for 10 mins, 100 nm nanodiamond: 2500 rpm for 10 mins) for 3 times to remove excessed neutravidin. Then, the nanodiamond-avidin conjugates were mixed with thiol-modified DNA (100 μM) for 2 hours at room temperature (300 rpm). The nanodiamond-DNA conjugates were purified by centrifugation (40 nm nanodiamond: 5000 rpm for 10 min, 100 nm nanodiamond: 2500 rpm for 10 min) for 3 times to remove excessed biotin-DNA.

#### DNA self-assembly of PEN nanoparticles

To construct the PEN nano-assembly, DNA-modified nanodiamonds (40 nm and 100 nm) were incubated with DNA-modified AuNPs (80 nm, 50 nm and 40 nm) in PBS solution with certain ratio for 30 min, and purified by centrifugation (1000 rpm for 2mins)

The following DNA sequences purchased from Integrated DNA Technologies were used in our work:

5’ ATGATATAGACGTTGTGGCAAAAAAAA-thiol 3’

5’ GCCACAACGTCTATATCATAAAAAAAA-biotin3’

#### Correlative imaging

The ITO glasses with a special fiducial marker on the surface were immersed in the piranha solution for 5 min followed by rinsing with copious amounts of ultrapure water. The cleaned ITO glasses were dried by N_2_ and then treated with oxygen plasma to make the surface hydrophilic (Plasma cleaner for 1 min at high radio frequency (RF) level). The purified PEN sample (100 µl) was left to adsorb on the surface of ITO glass for 10 min. Then, the ITO glasses were washed with ultrapure water and dried with N_2_ immediately. The sample was first scanned using SEM. We found the desired PEN particles near the laser marker under SEM image. Then, we relocated these particles under the dark-field microscopy and FLIM microscopy for separate studies.

#### SEM imaging

Tescan Mira 3 GMU field emission gun SEM with an acceleration voltage of 20 kV was used in this study.

#### Darkfield imaging

We use an upright microscope (Nikon, Japan) equipped with a dark-field condenser (numerical aperture (NA) = 1) and a 50× air objective lens (NA=0.8). The sample slides were immobilized on a platform, and a halogen lamp provided a white light source to excite the PEN particles to generate plasmon resonance scattering light. The scattered light was collected with a true-color digital camera to generate the dark-field color image and was also split with a monochromator (Kymera 328i, Andor, Oxford Instruments), and recorded with a spectrograph charge-coupled device (CCD) (iDus420, Andor, Oxford Instruments) to obtain the scattering spectra. The spectrum of an individual PEN particle was corrected by subtracting the background spectrum taken from the adjacent regions without the particle.

#### FLIM imaging

A customized confocal microscope (based on ISS Q2 laser scanning nanoscope) with single-molecule detection sensitivity was used for single particle intensity and fluorescence lifetime imaging. The excitation source was a Fianium supercontinuum SC-400 laser (6 psec pulse width and 42MHz repetition rate) equipped with a NKT acousto-optic tunable filter (AOTF) to select specific wavelength. Incident wavelength of 532 nm was used for exciting the PEN particles. The excitation light further cleaned-up by laser-line bandpass filter (532/10) was reflected by a dichroic mirror and focused onto the sample on an inverted microscope (Olympus, IX70). A water immersion objective (Olympus 60X, 1.2 numerical aperture (NA)) was used for focusing the laser light onto the sample and for collecting the fluorescence intensity emission from the sample. The fluorescence signal that passed through the dichroic mirror and a band-pass filter (650–720 nm, Chroma) was focused through a 75 μm pinhole to single photon avalanche photodiode (SPAD) (SPCM-AQRH-14, Excelitas Technologies) detector. The imaging in our set-up was performed with Galvo-controlled mirrors with related electronics and optics controlled through the 3X-DAC control card. The software module in ISS VistaVision for data acquisition and data processing along with this time-correlated single photon counting (TCSPC) module from Becker & Hickl (SPC-150) facilitate fluorescence lifetime measurements. The fluorescence intensity decays were analyzed in terms of an exponential model. The values of fluorescence lifetimes were determined using the Vistavision software with nonlinear least-squares fitting. Fluorescence lifetimes were estimated by fitting to a χ^2^value of less than 1.2 and a residual trace that was symmetrical about the zero axis. All the analyses were performed using Vistavision software and Originlab.

#### Cell culture and analysis

HeLa and GFP-HeLa cells were grown in Dulbecco’s modified Eagle’s medium (DMEM) (Millipore Sigma) culture medium with 10% fetal bovine serum (Millipore Sigma) and incubated at 37 °C, 5% CO_2_. HeLa cells were used because it was a widely studied cell line, and the results can provide useful information for those who are interested in using PEN nano-assemblies to probe cellular processes.

Confocal Imaging. Leica SP8 confocal was used for nanodiamond imaging with 532 nm laser excitation, and the emission channel was set to 650–700 nm. For GFP imaging, 488 nm laser was used for excitation, and the emission channel was set to 500–550 nm.

Cell Viability Assay. Cells were cultured in a 24-well microplate overnight and then incubated with PEN nanoparticles for 3 days. Next, the ethidium homodimer and Hoechst reagents (Sigma) was added and the cells were incubated in a humidified atmosphere for 15 mins. The microplate imaged with Nikon fluorescence microscopy, the blue channel is for Hoechst, and the red channel is for ethidium homodimer.

#### FDTD simulations

Ansys Lumerical 2020 R2.2 (Anasys. Inc. Vancouver, BC, Canada) software was used to perform all the FDTD simulations. A plane wave was used as the input light source for calculating the extinction spectra and the EM field and excitation rate enhancement. A mesh size of 1 nm was used for the former while a mesh size of 0.25 nm was used for the latter. An electric dipole oscillating horizontally was used as the input light source of emission rate enhancement with a mesh size of 1 nm. The intrinsic quantum efficiency for the electric dipole was modeled to be 0.7. Perfectly match layer (PML) boundary conditions were used for all the calculations. A refractive index of 2.42 was used to model the nanodiamond. The dielectric function for gold as from CRC.^44^ The background refractive index was set as 1.0.

## ASSOCIATED CONTENT

## NOTES

The authors declare no competing financial interest.

## ACKNOWLEDGMENT

This research was supported by National Institute of General Medical Sciences (DP2GM128198), National Institute of Biomedical Imaging and Bioengineering (2-P41-EB015871–31), AFOSR 21RT0264, and NIAID R01-AI150447.

## REFERENCE

1. Schirhagl, R., et al., Nitrogen-Vacancy Centers in Diamond: Nanoscale Sensors for Physics and Biology. Annual Review of Physical Chemistry 2014, 65 (1), 83–105.

2. Gruber, A., et al., Scanning Confocal Optical Microscopy and Magnetic Resonance on Single Defect Centers. Science 1997, 276 (5321), 2012–2014.

3. Taylor, J. M., et al., High-sensitivity diamond magnetometer with nanoscale resolution. Nature Physics 2008, 4 (10), 810–816.

4. Englund, D., et al., Deterministic Coupling of a Single Nitrogen Vacancy Center to a Photonic Crystal Cavity. Nano Letters 2010, 10 (10), 3922–3926.

5. Dolde, F., et al., Electric-field sensing using single diamond spins. Nature Physics 2011, 7 (6), 459–463.

6. Gong, J., et al., Nanodiamond-based nanostructures for coupling nitrogen-vacancy centres to metal nanoparticles and semiconductor quantum dots. Nature Communications 2016, 7 (1), 11820.

7. Giannini, V., et al., Plasmonic Nanoantennas: Fundamentals and Their Use in Controlling the Radiative Properties of Nanoemitters. Chemical Reviews 2011, 111 (6), 3888–3912.

8. Li, M., et al., Plasmon-enhanced optical sensors: a review. Analyst 2015, 140 (2), 386–406.

9. Willets, K. A.; Van Duyne, R. P., Localized Surface Plasmon Resonance Spectroscopy and Sensing. Annual Review of Physical Chemistry 2007, 58 (1), 267–297.

10. Jain, P. K., et al., On the Universal Scaling Behavior of the Distance Decay of Plasmon Coupling in Metal Nanoparticle Pairs: A Plasmon Ruler Equation. Nano Letters 2007, 7 (7), 2080–2088.

11. Gonçalves, P. A. D., et al., Plasmon–emitter interactions at the nanoscale. Nature Communications 2020, 11 (1), 366.

12. Kongsuwan, N., et al., Plasmonic Nanocavity Modes: From Near-Field to Far-Field Radiation. ACS Photonics 2020, 7 (2), 463–471.

13. Martín-Jiménez, A., et al., Unveiling the radiative local density of optical states of a plasmonic nanocavity by STM. Nature Communications 2020, 11 (1), 1021.

14. Shahbazyan, T. V., Local Density of States for Nanoplasmonics. Physical Review Letters 2016, 117 (20), 207401.

15. Sanders, S.; Manjavacas, A., Analysis of the Limits of the Local Density of Photonic States near Nanostructures. ACS Photonics 2018, 5 (6), 2437–2445.

16. Koya, A. N., et al., Novel Plasmonic Nanocavities for Optical Trapping-Assisted Biosensing Applications. Advanced Optical Materials 2020, 8 (7), 1901481.

17. Maccaferri, N., et al., Recent advances in plasmonic nanocavities for single-molecule spectroscopy. Nanoscale Advances 2021, 3 (3), 633–642.

18. Tang, J., et al., Selective far-field addressing of coupled quantum dots in a plasmonic nanocavity. Nature Communications 2018, 9 (1), 1705.

19. Choy, J. T., et al., Enhanced single-photon emission from a diamond–silver aperture. Nature Photonics 2011, 5 (12), 738–743.

20. Yu, H.-D., et al., Chemical routes to top-down nanofabrication. Chemical Society Reviews 2013, 42 (14), 6006–6018.

21. Schietinger, S., et al., Plasmon-Enhanced Single Photon Emission from a Nanoassembled Metal–Diamond Hybrid Structure at Room Temperature. Nano Letters 2009, 9 (4), 1694–1698.

22. Andersen, S. K. H., et al., Ultrabright Linearly Polarized Photon Generation from a Nitrogen Vacancy Center in a Nanocube Dimer Antenna. Nano Letters 2017, 17 (6), 3889–3895.

23. Bogdanov, S. I., et al., Ultrabright Room-Temperature Sub-Nanosecond Emission from Single Nitrogen-Vacancy Centers Coupled to Nanopatch Antennas. Nano Letters 2018, 18 (8), 4837–4844.

24. Rose, A., et al., Control of Radiative Processes Using Tunable Plasmonic Nanopatch Antennas. Nano Letters 2014, 14 (8), 4797–4802.

25. Hoang, T. B., et al., Ultrafast spontaneous emission source using plasmonic nanoantennas. Nature Communications 2015, 6 (1), 7788.

26. Li, C.-Y., et al., Observation of inhomogeneous plasmonic field distribution in a nanocavity. Nature Nanotechnology 2020, 15 (11), 922–926.

27. Kumar, S., et al., Overcoming evanescent field decay using 3D-tapered nanocavities for on-chip targeted molecular analysis. Nature Communications 2020, 11 (1), 2930.

28. Peng, Z.; Liu, H., Bottom-up Nanofabrication Using DNA Nanostructures. Chemistry of Materials 2016, 28 (4), 1012–1021.

29. Kuzyk, A., et al., Reconfigurable 3D plasmonic metamolecules. Nature Materials 2014, 13 (9), 862–866.

30. Kuzyk, A., et al., Selective control of reconfigurable chiral plasmonic metamolecules. Science Advances 3 (4), e1602803.

31. Sharma, A., et al., Chapter 3 - DNA nanostructures: chemistry, self-assembly, and applications. In Emerging Applications of Nanoparticles and Architecture Nanostructures, Barhoum, A.; Makhlouf, A. S. H., Eds. Elsevier: 2018; pp 71–94.

32. Fang, W., et al., Quantizing single-molecule surface-enhanced Raman scattering with DNA origami metamolecules. Science Advances 5 (9), eaau4506.

33. Li, Y., et al., Self-Assembly of Molecule-like Nanoparticle Clusters Directed by DNA Nanocages. Journal of the American Chemical Society 2015, 137 (13), 4320–4323.

34. Lermusiaux, L.; Funston, A. M., Plasmonic isomers via DNA-based self-assembly of gold nanoparticles. Nanoscale 2018, 10 (41), 19557–19567.

35. Chao, J., et al., DNA-based plasmonic nanostructures. Materials Today 2015, 18 (6), 326–335.

36. Barrow, S. J., et al., The surface plasmon modes of self-assembled gold nanocrystals. Nature Communications 2012, 3 (1), 1275.

37. Liang, L., et al., A Programmable DNA-Silicification-Based Nanocavity for Single-Molecule Plasmonic Sensing. Advanced Materials 2021, 33 (7), 2005133.

38. Fujisaku, T., et al., pH Nanosensor Using Electronic Spins in Diamond. ACS Nano 2019, 13 (10), 11726–11732.

39. Kucsko, G., et al., Nanometre-scale thermometry in a living cell. Nature 2013, 500 (7460), 54–58.

40. Wang, J., et al., High-sensitivity temperature sensing using an implanted single nitrogen-vacancy center array in diamond. Physical Review B 2015, 91 (15), 155404.

41. Pazos-Perez, N., et al., Modular assembly of plasmonic core–satellite structures as highly brilliant SERS-encoded nanoparticles. Nanoscale Advances 2019, 1 (1), 122–131.

42. Maiti, A., et al., Efficient Excitation of Higher Order Modes in the Plasmonic Response of Individual Concave Gold Nanocubes. The Journal of Physical Chemistry C 2017, 121 (1), 731–740.

43. Chen, J.-D., et al., Radiation of the high-order plasmonic modes of large gold nanospheres excited by surface plasmon polaritons. Nanoscale 2018, 10 (19), 9153–9163.

44. Lide, D. R., CRC handbook of chemistry and physics. Boca Raton, Fla. CRC Press/Taylor and Francis 2009, 2010, 486.

